# *AePUb* promoter length modulates gene expression in *Aedes aegypti*

**DOI:** 10.1101/2023.08.11.553001

**Authors:** Michelle A.E. Anderson, Philip T. Leftwich, Ray Wilson, Leonela Z. Carabajal Paladino, Sanjay Basu, Sara Rooney, Zach N. Adelman, Luke Alphey

## Abstract

Molecular tools for modulating transgene expression in *Aedes aegypti* are few. Here we demonstrate that adjustments to the *AePUb* promoter length can alter expression levels of two reporter proteins in *Ae. aegypti* cell culture and in mosquitoes. This provides a simple means for increasing or decreasing expression of a gene of interest and easy translation from cells to whole insects.

## Introduction

*Aedes aegypti* is a mosquito of medical importance to countries worldwide. This invasive pest has spread to every continent except Antarctica. It is the primary vector of the yellow fever virus, the dengue viruses, Zika virus and chikungunya virus, among others (1). These diseases cause the highest burden to tropical and subtropical areas and disproportionately affect the poorest populations. New technologies for the control of this invasive pest are required as widespread insecticide use has led to insecticide-resistant populations of this species.

Molecular tools are required to study this mosquito and develop new genetic strategies to control it. Most tools used today were originally developed in the model insect *Drosophila melanogaster*. The optimization of these for use in mosquitoes has enabled developments in gene editing tools such as CRISPR/Cas9 (2,3). Promoter fragments for the expression of genes of interest in both cell culture and whole insects play a crucial role in our ability to investigate this mosquito. There are a few select promoters identified that function in a wide range of tissues and cell types. Highly active *D. mel* promoters such as *DmAct5C* have been used (4). Other promoters such as Hr5/IE1 and OpIE2 are of of baculoviral origin (5,6) and were identified for use in *Drosophila* and then translated directly to mosquitoes. Relatively few *Ae. aegypti* native promoters have been characterized and used; exceptions include *UbL40* and *PUb (7)* and, more recently, *Hsp83 (8)*, which display ubiquitous expression. Th handful of promoters are used in various applications (9) and are frequently used to express mRNAs encoding fluorescent proteins, to provide markers for transgenesis/transfection, revealing the presence of a transgene construct otherwise lacking visible phenotype. Other promoters commonly characterized have tissue-specific expression patterns, such carboxypeptidase in the midgut, *zpg, nos, vasa* in ovaries or β2-*tubulin* in testes (10–13); this is useful for some genes of interest where expression in a specific tissue is vital. With advances in CRISPR/Cas9, new panels of germline specific promoters have also been characterized from *Ae. aegypti* (14,15).

A more refined set of promoters which modulate expression levels in a broad range of cell and tissue types would enable a more modular approach to research in *Ae. aegypti*. A single promoter that could be used in cultured cells and then directly used *in vivo* in insects could enable higher throughput screens that more easily translate from flask to insect. Expression of certain genes may prove detrimental or toxic to specific cells at high levels, and the ability to ‘de-tune’ expression would be advantageous. Here we sought to determine if the *PUb* promoter could be manipulated to enhance or decrease the expression of a reporter gene in both cells and transgenic *Ae. aegypti* mosquitoes.

## Materials and Methods

### Plasmids and cloning

Firefly and Renilla luciferase expression plasmids were cloned by standard methods starting with the pGL3 *PUb*-luc plasmid described previously (7) and pSLfa-*PUb*-*MCS* (Addgene plasmid # 52908). Transgenesis plasmids were generated using NEBuilder HiFi Assembly Master Mix (NEB) and primers listed in Supplementary Table 2. Complete sequences are available through NCBI accession numbers OR236189-OR236199 (16).

### Cells, transfections and luciferase assays

*Aedes aegypti* Aag2 cells, *Aedes albopictus* C6/36 and U4.4 cells were cultured as previously described (2). Briefly, cells were maintained at 28°C without CO_2_ or humidification. All cells were cultured in Leibovitz’s L-15 (Gibco) supplemented with 10% Fetal Bovine Serum (Gibco), 10% tryptose phosphate broth (Gibco) and 1% pen-strep (5,000 u/mL, Gibco). Cells were seeded into 96-well plates the day before transfecting with TransIT Pro (Mirus). Transfections were performed using 10ng/well of firefly expression plasmid and 5ng/well of *PUb*-RL Renilla luciferase normalization control plasmid (17). Two days after transfection cells were washed with phosphate buffered saline (PBS) and lysed in 50μl 1X passive lysis buffer. Luciferase assays were carried out as previously described with the Dual Luciferase Assay kit (Promega) and a GloMax+ plate reader (Promega).

### Analysis

We carried out all analyses in R version 4.1.0 (R Development Core Team). Data sets were summarised with the ‘tidyverse’ range of packages and figures were generated using ggplot2. Generalized linear mixed models were fitted with the glmmTMB package using a negative binomial distribution with a log-link function and summarized with emmeans (18, 19).

Briefly the FF/RR ratio was analysed with the promoter construct and cell lines as fixed factors with an interaction term. To account for the data structure, we included random effects for experimental replicate. Promoter length was considered as both a factorial and continuous variable with the best fit model found with a factorial design. Model residuals were checked for violations of assumptions with the DHARMa package (20). Pairwise contrasts were made with a tukey adjustment. The script is available on Github (https://github.com/Philip-Leftwich/AePUb-promoter-length-)

### Mosquitoes, transgenesis and rearing

*Aedes aegypti* were reared in insectary conditions with 27-28°C, 60-70% RH, and a 12/12 hour day/night cycle with one hour of dusk/dawn. Mosquitoes were provided 10% sucrose, *ad libitum*, and bloodfed on defibrinated horse blood (TCS) using a Hemotek artificial bloodfeeding system (Hemotek). All insect procedures were reviewed and approved by the Biological Agent and Genetic Modification Safety Committee (BAGMSC) at The Pirbright Institute.

Embryo microinjections were performed as previously described. Injection mixes comprised of 500ng/μl of *PUb* expression plasmid and 300ng/μl of AGG1733 Ae*PUb*(−565)⍰C31-SV40 3’UTR (21). The AGG1520 transgenic line which contains the 3xP3-mCherry-SV40 3’UTR transgenic marker, an attP docking site, and a secondary cassette not relevant to this study, was used for insertion of plasmids AGG2143-2146. This line has been identified by adapter-mediated PCR to be inserted on chromosome 2: 139436120-139437196 (reverse orientation) (unpublished). Insertion into the correct site was verified by PCR using the primers listed in Table 2.

### Imaging

Photographs of each life-cycle stage and dissected adult tissues (midgut and reproductive organs) were taken using a Leica M165FC fluorescence microscope fitted with an AmC filter. The magnification and exposure times were identical for each of the lines with respect to the life-cycle stage or tissue. Exposure times used were as follows: larvae 344ms; pupae 640ms; adult males and adult females 1500ms; male midguts 640ms; female midguts 485ms; testes 640ms and ovaries 485ms.

## Results

### *In vitro* expression in mosquito cells

The polyubiquitin (*PUb*, AAEL003888) derived promoter fragment is highly active during all life stages with constitutive expression in most tissues in *Aedes aegypti* mosquitoes. Initially characterised by Anderson et al (2010) this 1393 bp promoter fragment comprises 565bp of upstream sequence relative to the transcription start, then a transcribed region producing a 213bp 5’UTR after splicing removes a 615bp intron.

In total, we produced seven different variants of the *PUb* promoter, systematically increasing or decreasing the region upstream of the 5’UTR from -2500bp to ∼ 133bp (Fig 1). We also produced a version of this last promoter fragment (133bp), from which much of the intron was removed, retaining only the splice junctions and 41bp and 36bp of genomic sequence from the 5’ and 3’ of the intron respectively.

**Figure 1.**
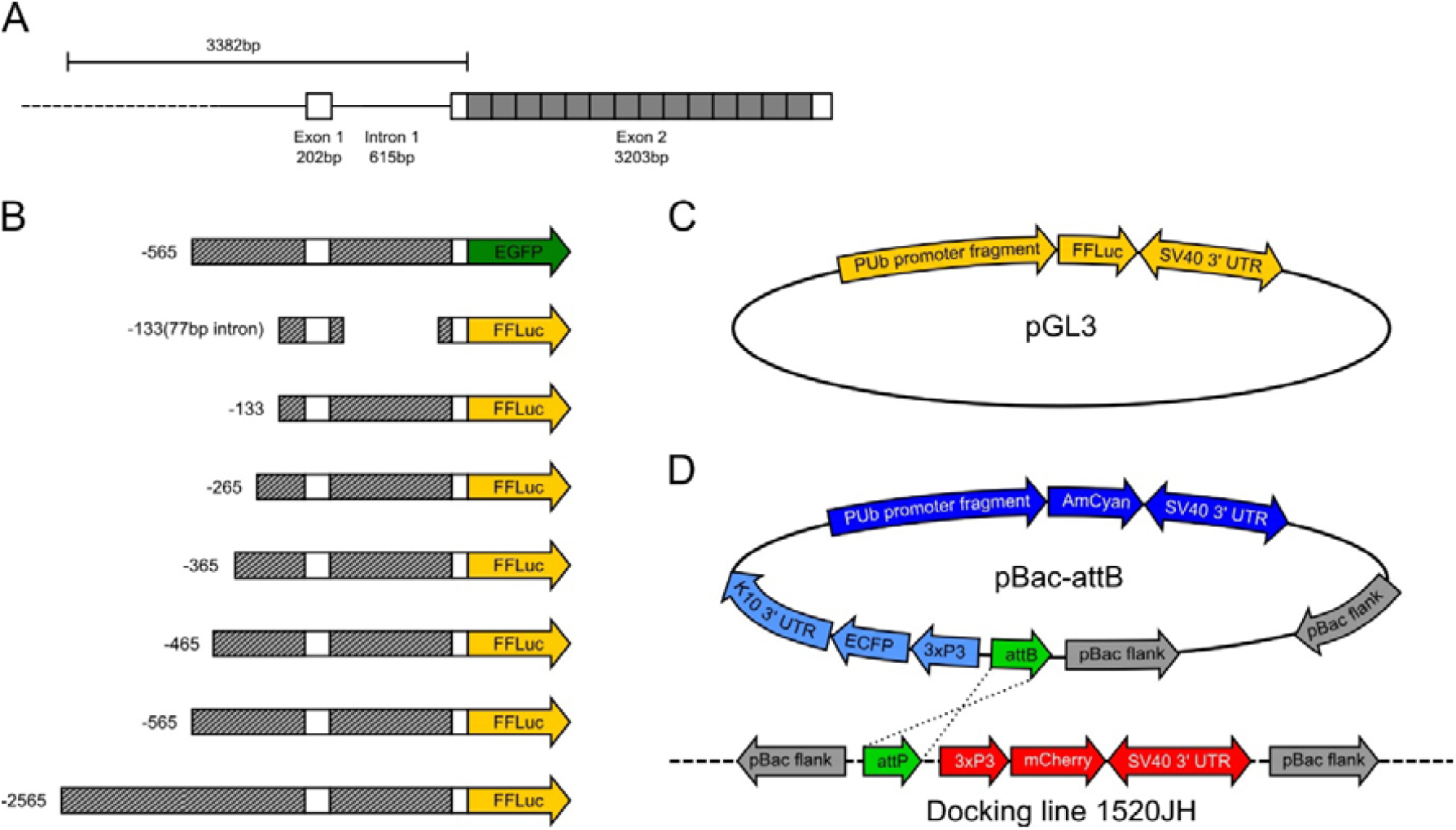
Representation of plasmid constructs. Diagram of *Aedes aegypti* AAEL003888 gene structure, adapted from Anderson et al 2010 (7). Promoter fragments are designated by the number of nucleotides upstream of the transcription start site (TSS=0). Solid grey boxes indicate ubiquitin monomers, white boxes indicate UTR (A). Diagram of putative promoter fragments cloned into reporter plasmids (B). Luciferase reporter plasmid used in cell culture experiments (C). AmCyan reporter plasmid and φC31 docking line used for transgenesis experiments (D).

We determined the transcriptional activity of all seven of these synthetic *PUb* promoter sequences by expressing a firefly luciferase (FF) gene in three cell lines derived from disease-relevant Culicine mosquitoes (*A. aegypti and A. albopictus*) using a previously described dual-luciferase assay (Fig 2).

**Figure 2.**
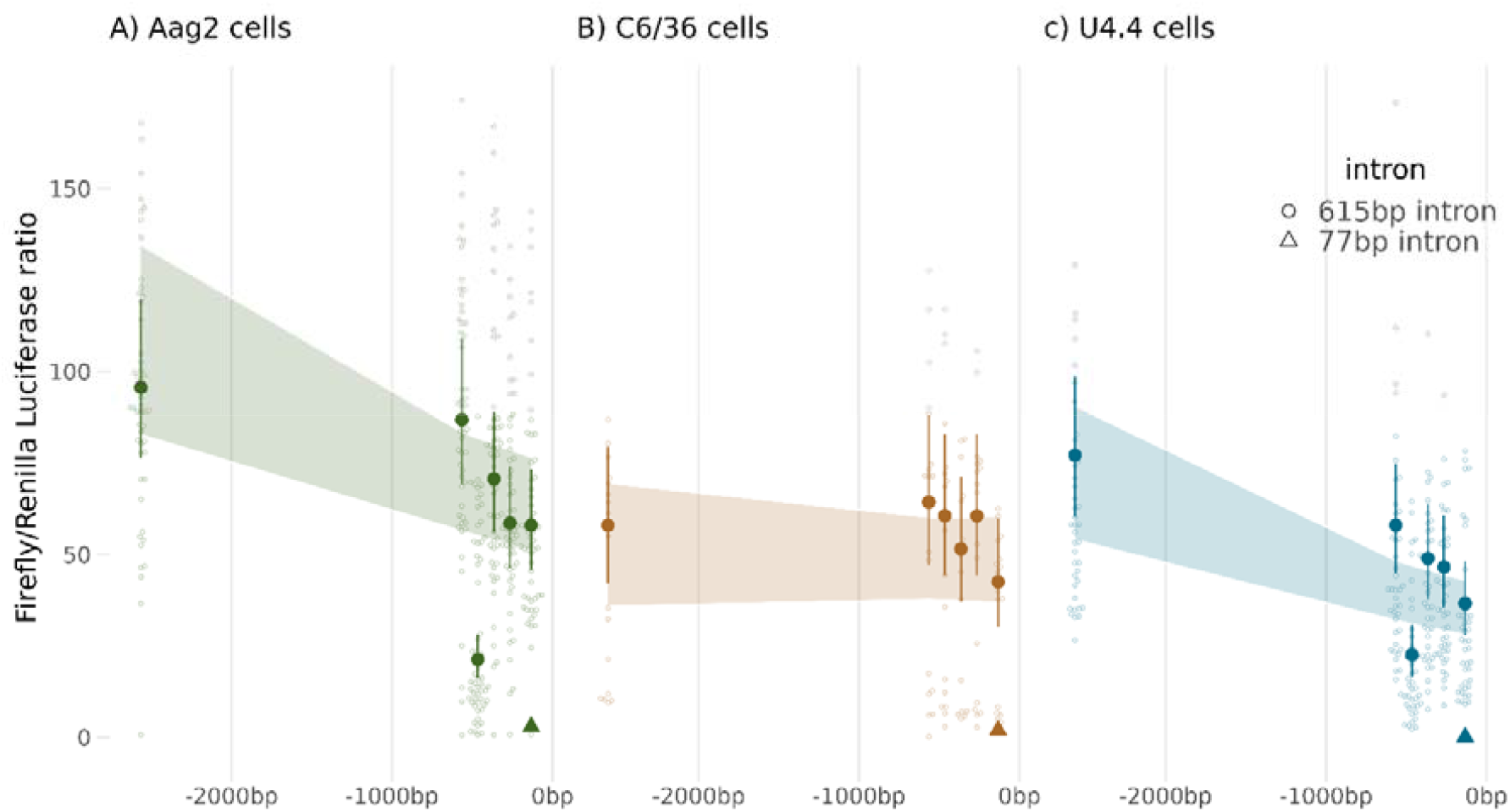
*PUb* promoter activity in vitro correlates with length. Ratios of FF/RL luciferase normalized to a GFP only control. Promoters are organized in order of distance (bp) 5’ of the transcriptional initiation start site (0bp). Large symbols and error bars (vertical lines) represent estimated mean and 95% confidence intervals for each promoter construct calculated by a generalized linear mixed model, with a negative binomial (‘log’ link) error distribution, with raw data shown as small symbols. Circles represent promoter sequences with a full-length intronic sequence, Triangles represent promoters with the truncated 77bp intronic sequence. Shaded areas represent the 95% confidence intervals for mean transcriptional activity modelled with length of promoter (bp) as a continuous variable.

We found a highly replicable pattern of gene expression across technical replicates, and levels of promoter activity were broadly in line with the species origin of the promoter, *PUb* activity in *U4*.*4* cells was only 81% [95% CI: ± 67-97%] and 61% [± 47-79%] in *C6/36* cells compared to *Aag2* cells (Table 1). Overall, there was limited evidence of differential responses in transcriptional activity to promoter editing between cell lines, indicating that the critical components of transcription in this promoter work in an essentially identical manner across species.

**Table 1.** Fixed and random effects table for the generalized linear mixed model (GLMM) fitted to the Luciferase Ratio detected in the engineered *PUb* promoter truncations.

| *Predictors* | **Values** |  |  |
| --- | --- | --- | --- |
|  | *Estimates* | *CI* | *p* |
| **(Intercept)** | 95.65 | 76.44 – 119.70 | **<0.001** |
| **Promoter-565** | 0.91 | 0.79 – 1.04 | 0.158 |
| **Promoter-465** | 0.22 | 0.18 – 0.27 | **<0.001** |
| **Promoter-365** | 0.74 | 0.64 – 0.85 | **<0.001** |
| **Promoter-265** | 0.61 | 0.53 – 0.71 | **<0.001** |
| **Promoter-133** | 0.61 | 0.52 – 0.70 | **<0.001** |
| **Promoter-133 (77bp intron)** | 0.03 | 0.02 – 0.04 | **<0.001** |
| **Promoter [no FF]** | 0.00 | 0.00 – Inf | 0.991 |
| **cell line [C636]** | 0.61 | 0.47 – 0.79 | **<0.001** |
| **cell line [u4.4]** | 0.81 | 0.67 – 0.97 | **0.022** |
| **Promoter-565:cell_lineC636** | 1.22 | 0.86 – 1.74 | 0.266 |
| **Promoter-465:cell_lineC636** | 4.68 | 3.18 – 6.89 | **<0.001** |
| **Promoter-365:cell_lineC636** | 1.20 | 0.83 – 1.74 | 0.327 |
| **Promoter-265:cell_lineC636** | 1.71 | 1.19 – 2.45 | **0.004** |
| **Promoter-133:cell_lineC636** | 1.21 | 0.82 – 1.78 | 0.333 |
| **Promoter-133 (77bp intron):cell_lineC636** | 1.13 | 0.46 – 2.78 | 0.796 |
| **Promoter [no FF] × cell line [C636]** | 0.70 | 0.00 – Inf | 1.000 |
| **Promoter-565:cell_lineu4.4** | 0.83 | 0.65 – 1.06 | 0.127 |
| **Promoter-465:cell_lineu4.4** | 1.31 | 0.95 – 1.82 | 0.102 |
| **Promoter-365:cell_lineu4.4** | 0.86 | 0.67 – 1.11 | 0.237 |
| **Promoter-265:cell_lineu4.4** | 0.99 | 0.76 – 1.28 | 0.914 |
| **Promoter-133:cell_lineu4.4** | 0.78 | 0.60 – 1.02 | 0.075 |
| **Promoter-133 (77bp intron):cell_lineu4.4** | 0.06 | 0.01 – 0.42 | **0.005** |
| **Promoter [no FF] × cell line [u4.4]** | 0.40 | 0.00 – Inf | 1.000 |
| **Random Effects** |  |  |  |
| **$\sigma^2$** | 0.27 |  |  |
| **$\tau_{00}$ experiment** | 0.10 |  |  |
| **ICC** | 0.28 |  |  |
| **N<sub>experiment</sub>** | 10 |  |  |
| **N<sub>obs</sub>** | 941 |  |  |
| **Observations** | 941 |  |  |
| **Marginal R<sup>2</sup> / Conditional R<sup>2</sup>** | 0.993 / 0.995 |  |  |

**Table 2.**
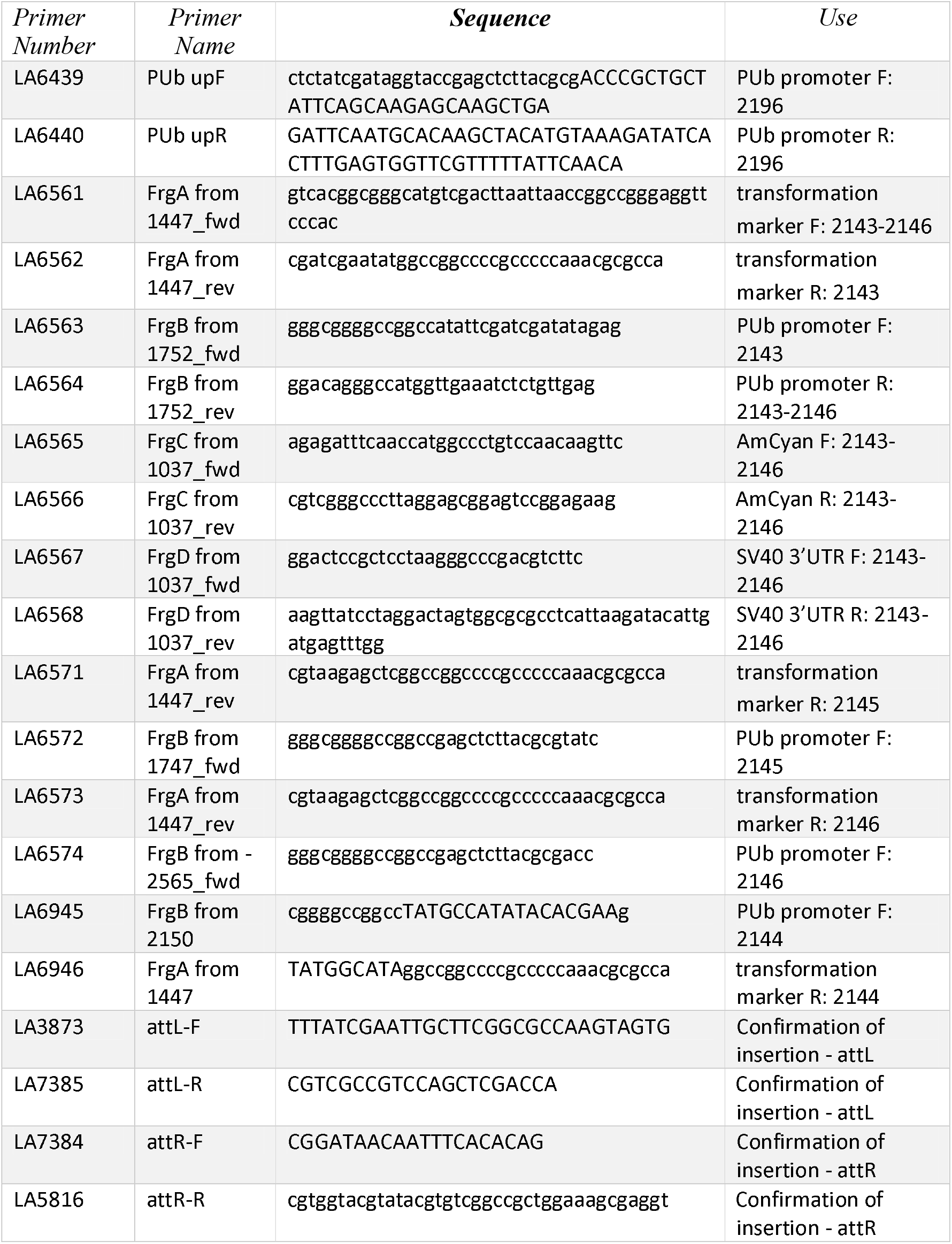
Primers used in this study.

Truncations of the promoter region produced an exponential drop in transcriptional activity of roughly 8% for every 500bp removed from the 5’ of the sequence, however, this model was not quite as good a model fit as comparing each promoter construct as an independent factor, and we observed a steeper drop in transcriptional activity in truncations closer to the transcription initiation site. This most likely indicates that transcription factor binding sites or other important regulators of transcriptional activity cluster within the 500bp 5’ of the transcription initiation site in this promoter.

The PUb(−133) promoter construct had only 61% [± 0.52-0.70] of the transcriptional activity of the full-length promoter -2565, and this fell to only 3% [± 0.02-0.04] in the -133(77bp intron) promoter sequence.

In Aag2 and U4.4 cells, we observed that by adjusting the length of the fragment upstream of the TSS we could modulate expression. In all cell lines the -133(77bp intron) was not significantly different from the no Firefly luciferase or -565 EGFP controls, and all other samples were significantly different from these three. This likely indicates that some positive regulatory elements are contained within the intron of the 5’UTR of this gene or that correct splicing has been disrupted. The pattern of modulation of expression by promoter length was not observed in C6/36 cells, where only intron removal produced a significant change in transgene expression in pairwise contrasts against other fragments.

### *In vivo* expression in *A. aegypti*

We selected four promoter fragments that were assessed *in vitro* for analysis *in vivo*. We selected the shortest fragment -133(77bp intron) with the lowest expression levels, an intermediate fragment -265, the previously published -565 fragment and the longest and highest expressing promoter fragment -2565 to express AmCyan from a transgene. It is well known that the genomic position of transgenes can influence expression levels. To avoid this “position effect” confounding comparison of different transgenic insertions, we used ⍰C31-mediated recombination to insert the experimental cassettes into a known, and previously characterised, insertion site which generated stable expression for previous constructs, AGG1520. This line contains a 3xP3-mCherry marker and an additional cassette irrelevant to this study.

The lines were generated by standard embryo microinjection of the donor plasmid and the ⍰C31 - helper and the insertions were confirmed by PCR. AmCyan fluorescence was imaged with standardized settings (Figs 3-4). No fluorescence could be detected in the –133(77bp intron) transgenics in any life stage or tissue. A small amount of fluorescence could be detected from –265 in the thorax of larvae, Malpighian tubules of male and female adults as well as the fore- and mid-gut of females. No expression was observed in the reproductive organs (Figs 4, 5 and S1) from this promoter fragment. As described previously, expression of AmCyan from the –565 promoter fragment could be readily observed in larvae and pupae, through the cuticle of adult males and females and in the gut of both male and female adults (Fig 3-4). In contrast to the previous publication characterizing this promoter (7) we did not observe substantial levels of expression in ovaries, even after a blood meal (Fig S1). This may be an indication that this genomic locus is somewhat less favourable for expression from this promoter than the originally characterized line where expression in ovaries was observed. We could also detect expression in the testes of adult males, more concentrated in the spermatozoa. A much more robust expression could be observed with the –2565 promoter across all stages and tissues.

**Figure 3.**
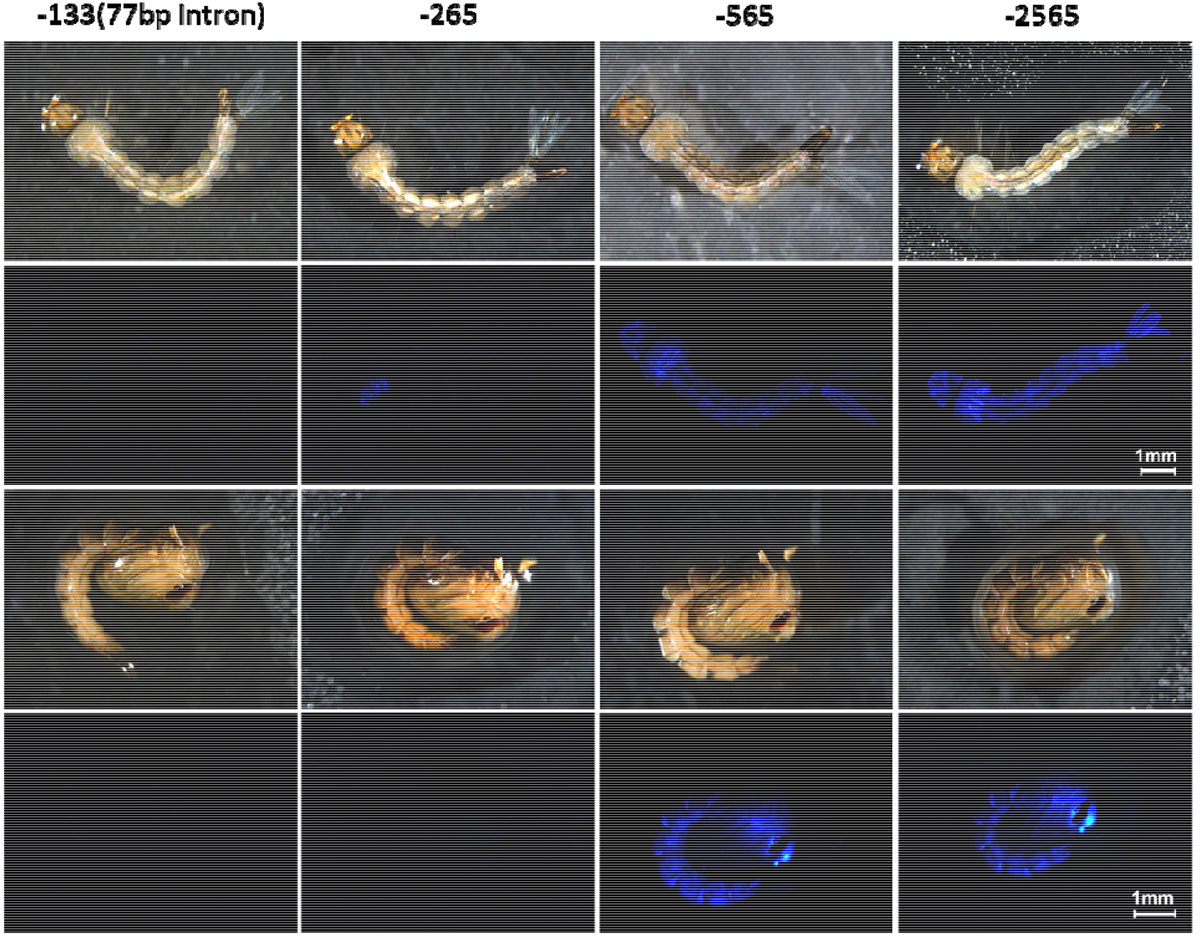
*PUb* promoter expression across developmental stages in transgenic *A. aegypti*. Brightfield and AmCyan fluorescence images of larvae (top two rows) and pupae (bottom two rows) with four different *PUb* promoter lengths (number indicates bp upstream of the transcription start site).

**Figure 4.**
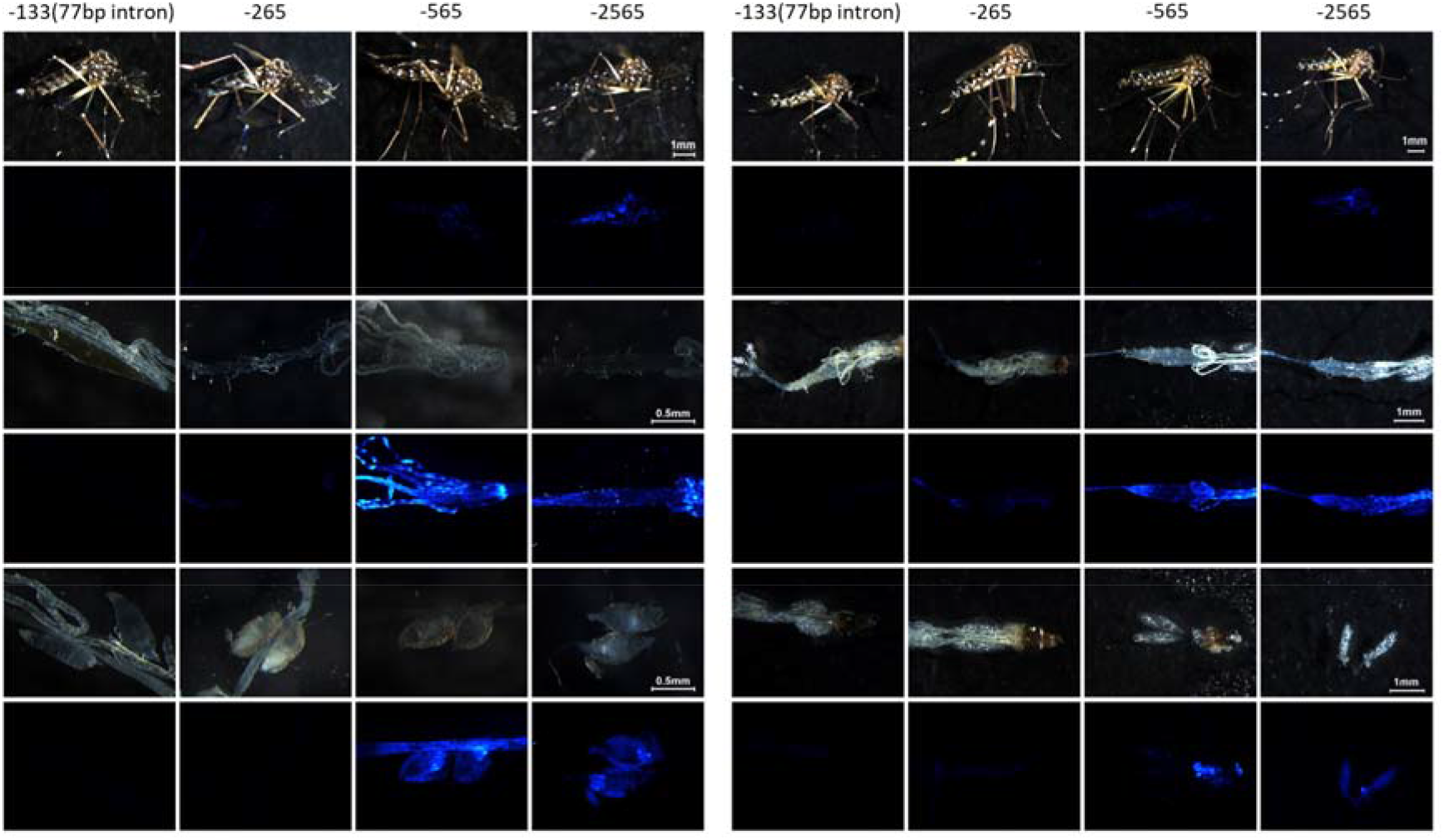
*PUb* promoter expression across adult tissues in transgenic *A. aegypti*. Brightfield and AmCyan fluorescence for adult males, dissected gut, and testes (left panels, top, middle and bottom, respectively). Brightfield and AmCyan fluorescence for adult females, dissected gut, and ovaries (right panels, top, middle and bottom, respectively).

## Discussion

This study investigated the transcriptional activity of *polyubiquitin* (*PUb*) promoter sequences in Culicine mosquitoes and cell lines. Our findings provide insights into the functional properties of the *PUb* promoter and shed light on the importance of specific regions, namely the intron within the 5’UTR, for gene expression.

One of the key findings of our study is the consistent pattern of gene expression observed across technical replicates. The observed levels of promoter activity were broadly in line with the species origin of the promoter, with the highest activity in Aag2 cells compared to C6/36 and U4.4 cells, suggesting that fundamental mechanisms of transcriptional regulation in the *PUb* promoter are largely conserved across these mosquito species. Truncations of the promoter fragment produced a roughly exponential decline in gene activity, with a severe decline in activity with a truncated intronic sequence. This abrupt decline indicates that some important sequences that regulate expression may be situated within the intron rather than 5’ to the transcription start.

Our *in vivo* work used a ⍰C31-mediated recombination technique to provide a fixed genomic integration site, allowing us to study the effects of promoter manipulation without the noise of random genomic integration sites. Consistent with the cell culture data, PUb-133 (77bp intron) expression of an AmCyan fluorescent marker was undetectable in our samples or tissues. At the same time, expression from promoters with intact intronic sequences was increasingly bright and ubiquitous as promoter fragment length increased. Interestingly, the full-length promoter sequence produced both the brightest fluorescence and the broadest tissue expression, while -565 and -265 showed increasingly dimmer and tissue-restricted expression. This may indicate that the loss of elements can include enhancers, silencers, or binding sites for transcription factors required for proper regulation of gene expression, with the absence of these regulatory elements in the shorter fragment leading to tissue-specific variation in visibility. It is also possible that the -565 fragment is more susceptible to the influence of neighbouring chromatin, while the -2565 fragment is better insulated from this. A wealth of future work is available to elucidate the relative importance of genomic insertion effects, tissue-specific effects, intron-based gene regulation and potential insulators of transgene expression.

Our study provides valuable insights into the transcriptional activity of synthetic *PUb* promoter fragments in *A. aegypti* mosquitoes. Characterizing these promoter fragments and identifying genomic locus influences contribute to expanding the genetic toolbox for precise gene expression manipulation in *A. aegypti*, facilitating further investigations into mosquito biology and the development of targeted vector control strategies.

## Supporting information

Supplemental materia S1

## Acknowledgements

MAEA was funded by Defense Advanced Research Projects Agency (DARPA) award [N66001-17-2-4054] to Kevin Esvelt at MIT. PTL, RW and LZCP were funded by a Wellcome Trust Investigator Award [110117/Z/15/Z] to LA. LA was supported through strategic funding from the UK Biotechnology and Biological Sciences Research Council (BBSRC) to The Pirbright Institute (BBS/E/I/00007033, BBS/E/I/00007038 and BBS/E/I/00007039). The views, opinions and/or findings expressed are those of the authors and should not be interpreted as representing the official views or policies of the U.S. Government. The funders had no role in study design, data collection and analysis, decision to publish, or preparation of the manuscript.

## Author contributions

MAEA, LZCP and RW performed the experiments. PTL analysed the data. MAEA, ZNA and LA conceived the experiments. SB and SR provided reagents. MAEA and PTL wrote the first draft of the manuscript and all authors reviewed and approved the manuscript.

## Data availability statement

All data generated is included in the manuscript and supplemental files.

## Competing interest statement

The authors declare they have no competing interests.

## Notes

### Competing Interest Statement

The authors have declared no competing interest.

## References

1. Viglietta M, Bellone R, Blisnick AA, Failloux AB. Vector Specificity of Arbovirus Transmission. Front Microbiol [Internet]. 2021 [cited 2023 Jun 17];12. Available from: https://www.frontiersin.org/articles/10.3389/fmicb.2021.773211

2. Anderson MAE, Purcell J, Verkuijl SAN, Norman VC, Leftwich PT, Harvey-Samuel T, et al. Expanding the CRISPR Toolbox in Culicine Mosquitoes: In Vitro Validation of Pol III Promoters. ACS Synth Biol. 2020 Mar 20;9(3):678–81.

3. Rozen-Gagnon K, Yi S, Jacobson E, Novack S, Rice CM. A selectable, plasmid-based system to generate CRISPR/Cas9 gene edited and knock-in mosquito cell lines. Sci Rep. 2021 Jan 12;11:736.

4. Pinkerton AC, Michel K, O’Brochta DA, Atkinson PW. Green fluorescent protein as a genetic marker in transgenic Aedes aegypti. Insect Mol Biol. 2000 Feb;9(1):1–10.

5. Pfeifer TA, Hegedus DD, Grigliatti TA, Theilmann DA. Baculovirus immediate-early promoter-mediated expression of the Zeocin resistance gene for use as a dominant selectable marker in dipteran and lepidopteran insect cell lines. Gene. 1997 Apr 1;188(2):183–90.

6. Theilmann DA, Stewart S. Molecular analysis of the trans-activating IE-2 gene of Orgyia pseudotsugata multicapsid nuclear polyhedrosis virus. Virology. 1992 Mar;187(1):84–96.

7. Anderson M a. E, Gross TL, Myles KM, Adelman ZN. Validation of novel promoter sequences derived from two endogenous ubiquitin genes in transgenic Aedes aegypti. Insect Mol Biol. 2010;19(4):441–9.

8. Webster SH, Scott MJ. The Aedes aegypti (Diptera: Culicidae) hsp83 Gene Promoter Drives Strong Ubiquitous DsRed and ZsGreen Marker Expression in Transgenic Mosquitoes. J Med Entomol. 2021 Nov 9;58(6):2533–7.

9. Biomolecules | Free Full-Text | Use of Insect Promoters in Genetic Engineering to Control Mosquito-Borne Diseases [Internet]. [cited 2023 Jun 17]. Available from: https://www.mdpi.com/2218-273X/13/1/16

10. Kyrou K, Hammond AM, Galizi R, Kranjc N, Burt A, Beaghton AK, et al. A CRISPR–Cas9 gene drive targeting doublesex causes complete population suppression in caged Anopheles gambiae mosquitoes. Nat Biotechnol. 2018 Nov;36(11):1062–6.

11. Adelman ZN, Jasinskiene N, Onal S, Juhn J, Ashikyan A, Salampessy M, et al. nanos gene control DNA mediates developmentally regulated transposition in the yellow fever mosquito Aedes aegypti. Proc Natl Acad Sci. 2007 Jun 12;104(24):9970–5.

12. Gantz VM, Jasinskiene N, Tatarenkova O, Fazekas A, Macias VM, Bier E, et al. Highly efficient Cas9-mediated gene drive for population modification of the malaria vector mosquito Anopheles stephensi. Proc Natl Acad Sci. 2015 Dec 8;112(49):E6736–43.

13. Smith RC, Walter MF, Hice RH, O’Brochta DA, Atkinson PW. Testis-specific expression of the β2 tubulin promoter of Aedes aegypti and its application as a genetic sex-separation marker. Insect Mol Biol. 2007;16(1):61–71.

14. Anderson MAE, Gonzalez E, Ang JXD, Shackleford L, Nevard K, Verkuijl SAN, et al. Closing the gap to effective gene drive in Aedes aegypti by exploiting germline regulatory elements. Nat Commun. 2023 Jan 20;14(1):338.

15. Li M, Bui M, Yang T, Bowman CS, White BJ, Akbari OS. Germline Cas9 expression yields highly efficient genome engineering in a major worldwide disease vector, Aedes aegypti. Proc Natl Acad Sci. 2017 Dec 5;114(49):E10540–9.

16. Lehwark P, Greiner S. GB2sequin - A file converter preparing custom GenBank files for database submission. Genomics. 2019 Jul 1;111(4):759–61.

17. Aryan A, Anderson MAE, Myles KM, Adelman ZN. Germline excision of transgenes in Aedes aegypti by homing endonucleases. Sci Rep. 2013 Apr 3;3(1):1603.

18. Brooks M E, Kristensen K, Benthem K J, van Magnusson A, Berg C W, Nielsen A, et al. glmmTMB Balances Speed and Flexibility Among Packages for Zero-inflated Generalized Linear Mixed Modeling. R J. 2017;9(2):378.

19. Lenth RV. emmeans: Estimated Marginal Means, aka Least-Squares Means [Internet]. 2023. Available from: https://github.com/rvlenth/emmeans

20. Hartig F. DHARMa: residual diagnostics for hierarchical (multi-level/mixed) regression models [Internet]. 2022 [cited 2023 Jun 17]. Available from: https://cran.r-project.org/web/packages/DHARMa/vignettes/DHARMa.html

21. Paladino LZC, Wilson R, Tng PY, Dhokiya V, Keen E, Cuber P, et al. Optimizing CRE and PhiC31 mediated recombination in Aedes aegypti [Internet]. bioRxiv; 2023 [cited 2023 Jul 11]. p. 2023.07.07.548128. Available from: https://www.biorxiv.org/content/10.1101/2023.07.07.548128v1

