## Supplemental materia S1 for "*AePUb* promoter length modulates gene expression in *Aedes aegypti*"

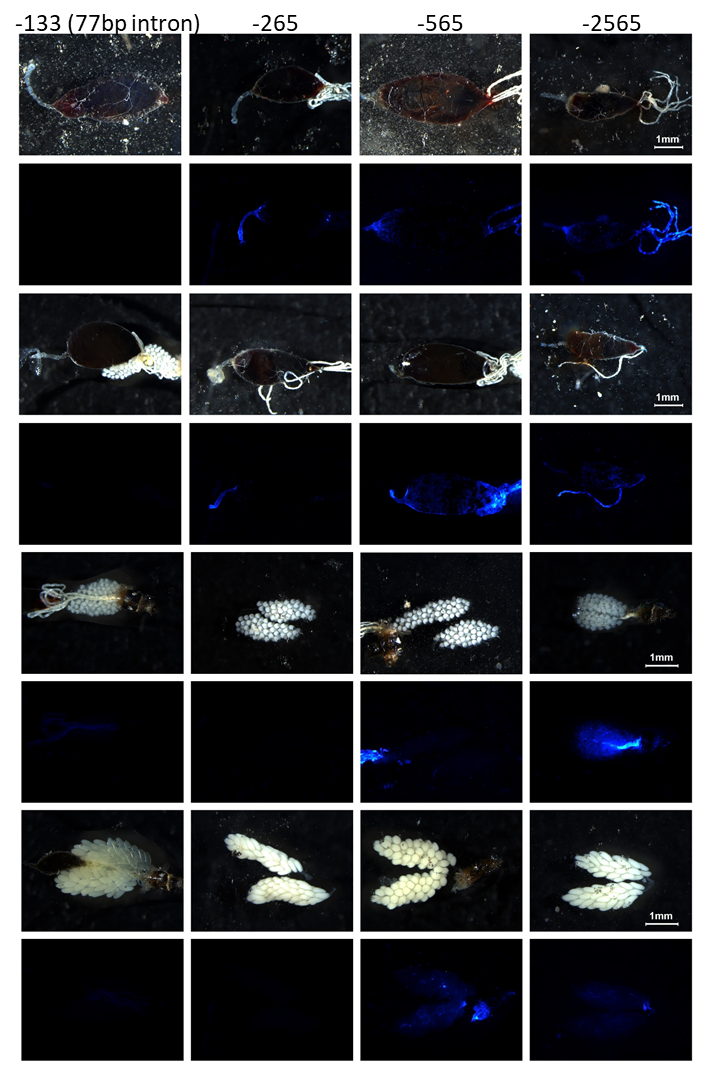

Figure S1. PUb promoter fragments do not express in post-blood meal midguts or ovaries. Brightfield and AmCyan fluorescence of 24h (top) and 48h (second set of panels) post bloodmeal midguts. Brightfield and AmCyan fluorescence of 24h (third set of panels) and 48h (bottom panels) post bloodmeal ovaries.
